# Model for FRET sensors of allosteric switchable modulator proteins for the determination of allosteric parameters

**DOI:** 10.1101/2022.08.20.503106

**Authors:** Heiko Babel

## Abstract

FRET-sensors are a well-established method to investigate protein-protein interactions. To determine how FRET-sensor can be employed for the study of switchable allosteric modulator proteins (SAMPs) I extend a previously established model for enzymatic SAMPs to include a FRET-sensor system. Using this model, I determine the prerequisites for using FRET to investigate modulator-regulator interaction. The model shows, that under saturating stimulus conditions only a trimolecular complex contributes to the measured FRET value. How the signal is relayed by the modulator can be investigated by comparing FRET values of unstimulated and signal-saturated sensor systems. Finally, to determine the allosteric mode of signal transduction the natural logarithm of the ratio of stimulated and unstimulated FRET efficiencies is a useful metric.

## 1 Introduction

Förster Resonance Energy Transfer (FRET) has been established in recent years as a fruitful method to investigate protein-protein interactions[1]. FRET relies on the nonradiative energy transfer between an excited donor-fluorophor and an acceptor fluorophor, which finally leads to fluorescence emission of the acceptor fluorophor and reduces fluorescence of the donor-fluorophor [2].

The basis for the energy transfer lies in the spectral properties of the fluorophors. It is necessary that the emission spectrum of the donor-fluorophor overlaps with the excitation spectrum of the acceptor fluorophor. Because the energy transfer relies on dipole-dipole interaction, the fluorophors need to be located in close proximity and be oriented according to the dipole-fields. The rate of energy-transfer *k* is proportional to the squared interaction energy *U* between donor- and acceptor-fluorophor [2],

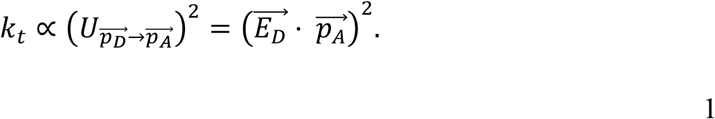

Here 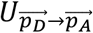 is the interaction-energy between donor- and acceptor dipole, 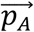 and 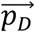 acceptor and donor-dipole respectively and 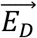 the field of the donor-dipole. How the acceptor-dipole needs to be aligned in the field of the donor-dipole is shown schematically in figure 1 A. Energy-transfer is maximal when the acceptor-dipole is aligned in a parallel fashion to the donor-field. Usually dipole-orientation and spectral properties are summarized in the factor *R*_*0*_^*6*^ which shows the relation of the FRET-efficiency to the donor-acceptor distance *r*_*DA*_:

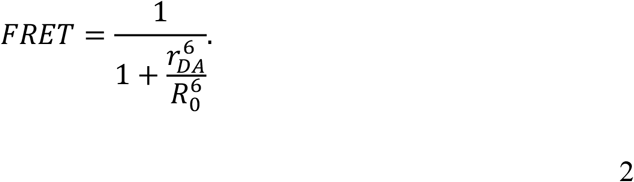

**Figure 1:**
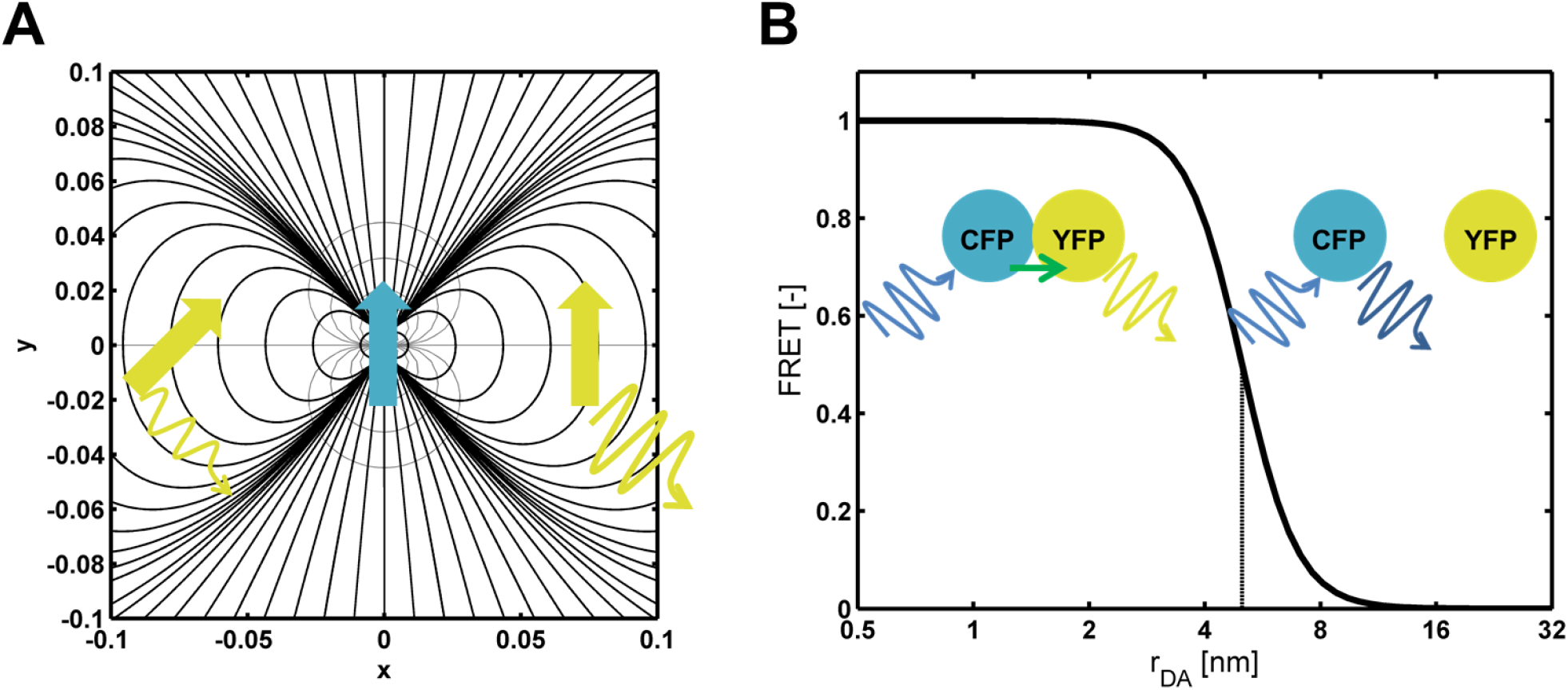
**Orientation and distance of fluorophores influence FRET efficiency. (A) Orientation of the acceptor-fluorophor-dipole in the donor-dipole field. Black lines show field-lines of the electric field, gray lines represent equipotential line of the electric potential. (B) Relationship between energy transfer efficiency and donor-acceptor distance.**

The influence of the distance on the efficiency of energy transfer is shown in figure 1 B. If the distance is larger than *R*_*0*_^*6*^ the rate of energy transfer decreases rapidly. If the fluorophores are not in a complex, no FRET can be observed.

To investigate protein-protein interactions the target proteins are labeled with a donor and acceptor fluorophore [3,4]. Due to the strong dependency on the distance, FRET can only occur *in vivo* when proteins form a complex. In addition, FRET measurements are dependent on the conformation of the proteins in the complex, since already small changes in distance or orientation of the fluorophores can influence the FRET-efficiency. This can be utilized for unimolecular FRET-sensor where a target protein is labeled with two fluorophors. Changes in protein conformation are then translated into changes in fluorophore orientation and finally influence FRET-efficiency [5].

Since the energy transfer increases fluorescence emission of the acceptor fluorophore this can be used to determine FRET. Since usually emission spectra of donor and acceptor fluorophores overlap, this method requires measurement of a set of controls to finally calculate the FRET efficiency. It is possible to obtain precise determination of FRET-efficiencies, but due to the control measurements it is a laborious method [6,7].

If a fluorophore is excited with the appropriate wavelength this excitation results in the emission of light. The exited state can also be relieved by release of thermal energy, fluorescence-quenching, by a photochemical reaction or bleaching of the fluorophor. FRET adds an additional pathway of relieve of the excited state. Consequently, FRET changes the effective rate by which fluorescence emission or photobleaching of the donor fluorophor occur. Both processes can be used to determine the FRET-efficiency [2,7].

In those cases the donor-fluorescence intensity or the photobleaching rate of the donor without the acceptor are measured. If the fluorescence intensity of the donor is measured the FRET-efficincy is given by:

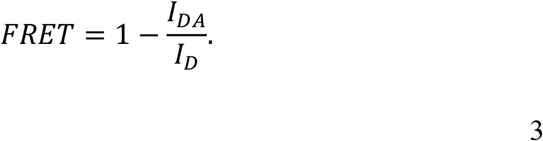

wherein *I*_*DA*_ and *I*_*D*_ are the donor-fluorescence-emission with and without the acceptor fluorophore respectively [2]. For this method either two samples, one containing the donor-fluorophore and one containing both donor and acceptor-fluorophore can be measured, or only a sample containing both donor- and acceptor-fluorophore are measured and the acceptor-fluorophore is inactivated. This can be achieved by acceptor-photobleaching. Here, first the sample with acceptor- and donor-fluorophore is measured, than the acceptor is specifically bleached with a laser and finally the donor-fluorescence is measured again. If the acceptor is bleached completely the FRET-efficiency can directly be calculated.

In this case the measured FRET-value depends on the properties of the protein complex, i.e. how the fluorophores are oriented in this complex, but also on the concentration of the proteins that are part of the complex. Since in the acceptor photobleaching method the donor-emission is measured, unbound donor molecules decrease the measured FRET-efficiency. The correct FRET-efficincy is than given by

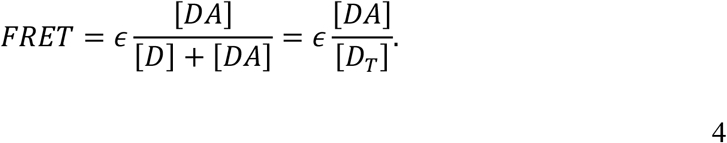

wherein ε is the efficiency of the energy transfer in the protein-complex, [*DA*] the concentration of the protein complex of fluorescently labeled proteins and [*D*_*T*_] the total concentration of donor-fluorophore labeled protein [7]. Theoretically the complex could be present in different conformations with different molecular FRET-efficiencies ε_i_, than the FRET-efficiency is given by:

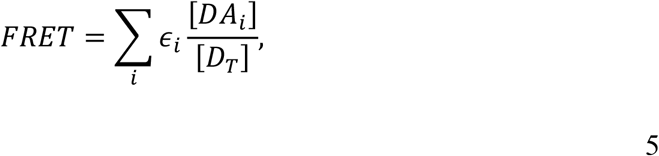

wherein [*DA*_*i*_] is the concentration of the *i*-th complex conformation.

Allosteric regulation is one of the key mechanisms for cellular signal transduction [8,9]. Although Rap-proteins are known for decades [10] only recently a series of structural studies showed that indeed Rap proteins are allosteric proteins which are monomers in solution and interact with a regulator protein in a 1:1 ratio [11–17]. Rap-proteins have one allosteric binding-site for the peptide, which is different from the large class of oligomeric protein receptors [18]. Of this protein family, the RapA modular belongs to the special class of enzymatic switchable allosteric modulator proteins (SAMPs) [19]. The RapA modulator dephorsphorylates the Spo0F regulator, thereby inactivating it. Its phosphatase activity is in turn allosterically regulated by the PhrA peptide [12,20]. A theoretical study showed, that an enzymatic SAMP such as RapA has two allosteric modes of signal transduction, the enzymatic allostery and so-called binding allostery. Enzymatic allostery is the molecular influence of the signal on the enzymatic activity of the modulator and binding allostery influence the affinity of the modulator to the regulator [19]. Recently, a FRET based biosensor system has been established to study the signal-transduction *in vivo* [21] and successfully been used to characterize the temporal signal transduction properties. The question remains if such a FRET system could be used to discern allosteric modes of an enzymatic SAMP *in vivo*.

In this study I extend the previously published enzymatic SAMP model to include a quantitative model of FRET measurements. With the help of this model the prerequisites for the detection of a FRET signal should be elucidated. I evaluate how signal stimulation influences the FRET response and how the FRET response depends on the molecular properties of the SAMP. Finally, experimental conditions are identified that provide insight into the *in vivo* allosteric mode of signal transduction.

## 2 Methods and Model

All numerical calculations were performed with Matlab R2011b ® Mathworks ® (Natick MA, USA). To calculate the steady-state of the ODE system, the ODE was solved numerically for a large time-point and the steady-state solution was verified using the fsolve-function. Analytical results were checked using Mathematica 8 of Wolfra Research (Champaign IL, USA).

### 2.1 FRET Signal of a SAMP System

When the donor-fluorophore is linked to the switchable allosteric modulator protein and the acceptor fluorophore is linked to the acceptor-fluorophore then the FRET efficiency for acceptor bleaching is given by:

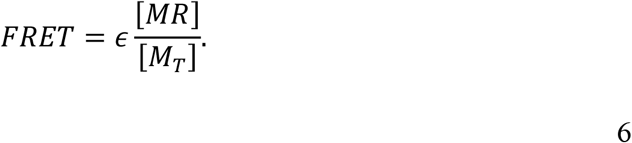

If the affinity to the modulator is strongly decreased by signal binding, the regulator can be released from the regulator modulator complex (Figure 3, upper path). In this case the signal correlates directly with the fraction of bound regulator. The signal can also not affect regulator affinity and only affect the enzymatic activity of the modulator. In this case, a complex of modulator, regulator and signal would be formed (Figure 3, lower path), which could in principle have the same concentration as the modulator-regulator complex. The trimolecular complex could have a different fluorophore orientation than the bimolecular modulator-regulator complex, so that the molecular FRET efficiency ε is altered. Both the bimolecular and trimolecular complex therefore contribute to the FRET signal:

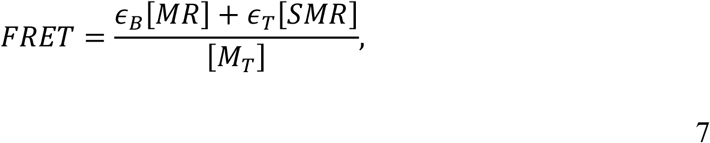

wherein ε_B_ and ε_*T*_ are the molecular FRET efficiency of the modulator-regulator complex and the trimolecular complex respectively.

**Figure 2:**
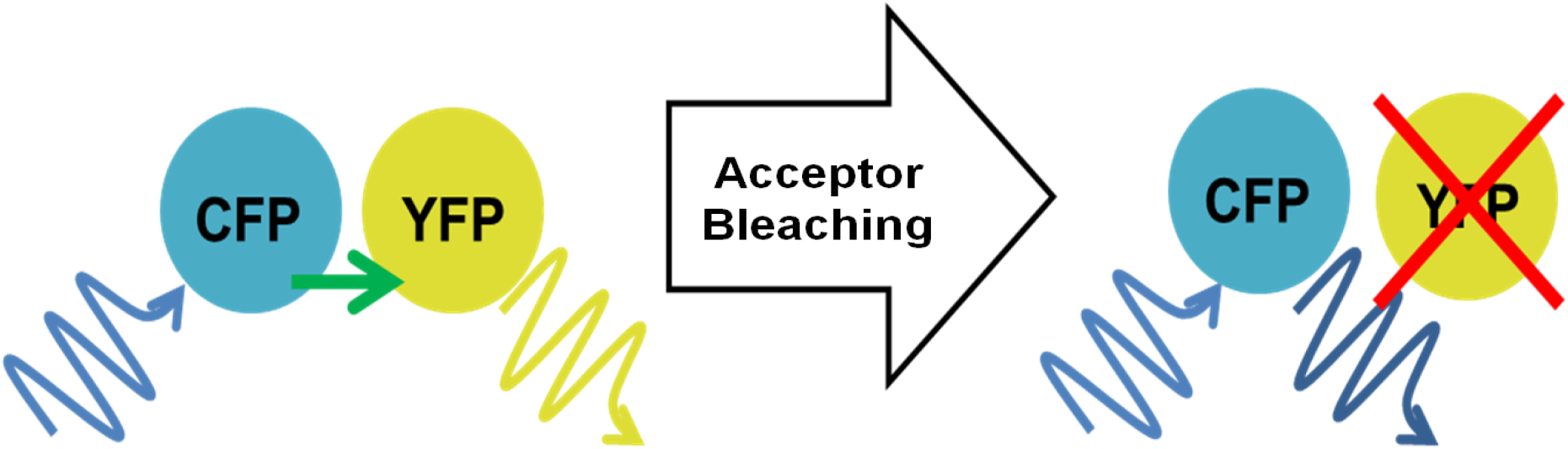
**Acceptorphotobleaching to determine the FRET-efficiency. The photoacceptor YFP is bleached specifically by a laser. This abolishes the FRET-energy tansfer and consequently increases fluorescence emission of the donor fluorophore CFP.**

**Figure 3:**
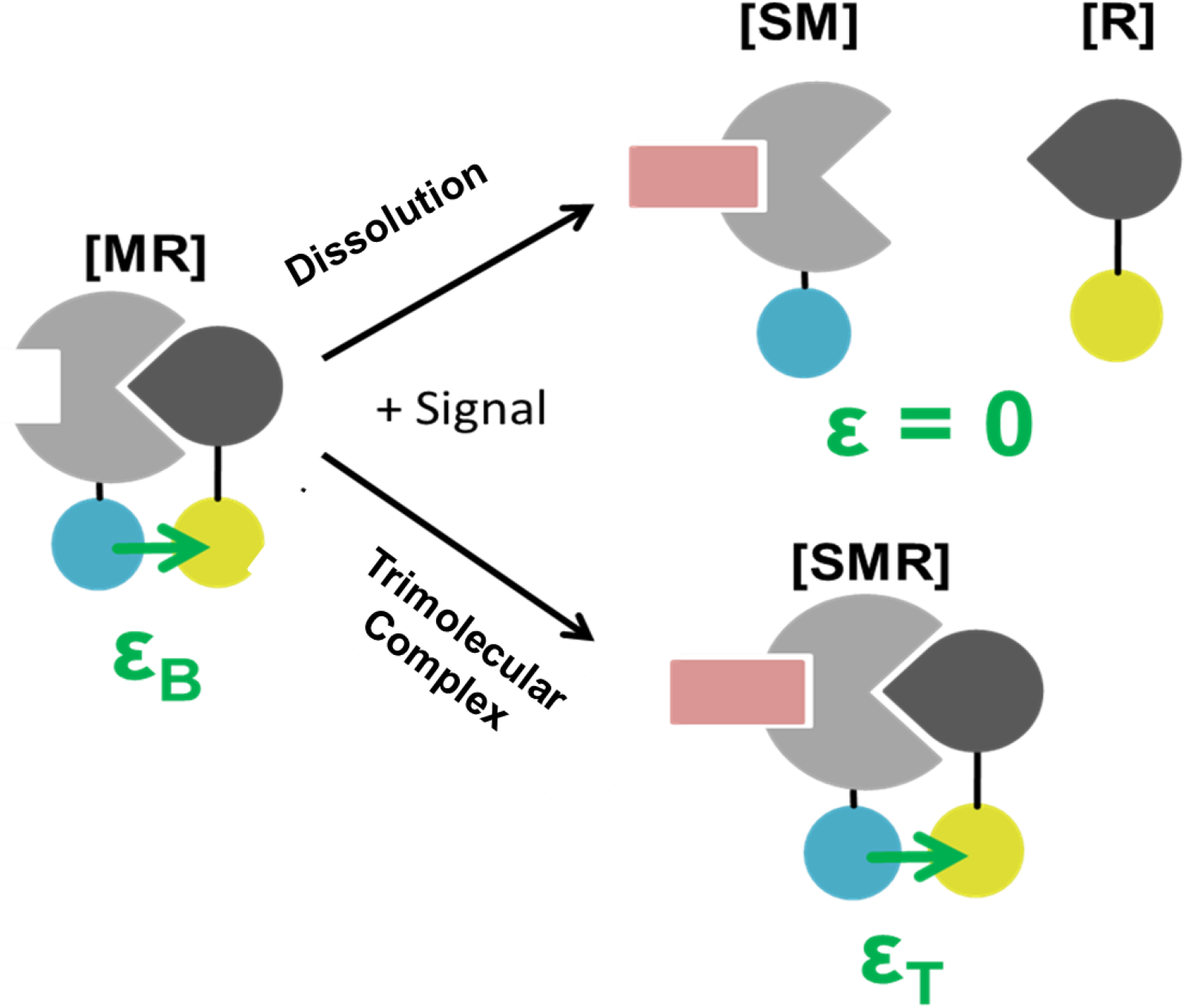
**Quantitiative model for bimolecular Modulator-Regulator sensor systems. Stimulation can lead to the formation of a trimolecular complex SMR with efficiency ε_*T*_ or the bimolecular MR complex with molecular FREt efficiency ε_B_ can be dissoluted by the addition of the signal.**

To understand how the FRET signal is affected by different allosteric modes I introduce a model for the interaction of modulator, regulator and signal.

### 2.2 Enzymatic SAMP-Model for FRET sensors

I use the model of the enzymatic SAMP as described previously [19] given by:

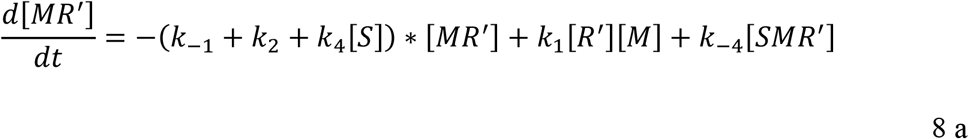

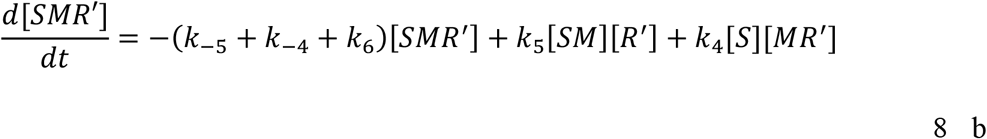

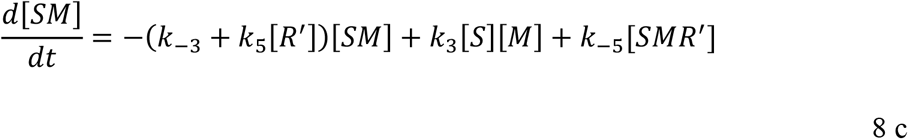

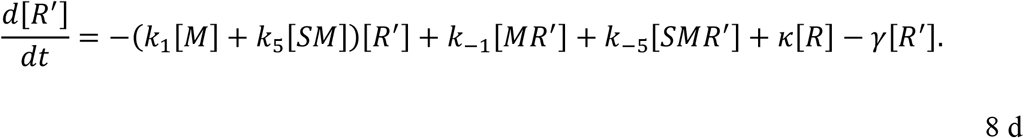

In brief, the active regulator R’, the modulator M and the signal S can form different complexes through reversible reactions. The modulator can inactivate the active regulator R’ with the rate k_2_ and in the case that the signal S is bound with rate k_6_. The inactive regulator R is activated with the rate κ and unspecifically inactivated with rate γ.

In addition, the total concentrations [R_T_], [S_T_] and [M_T_] are assumed to be constant:

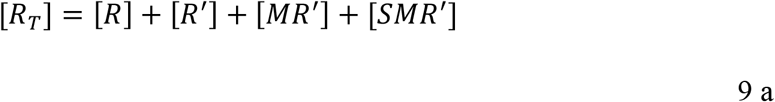

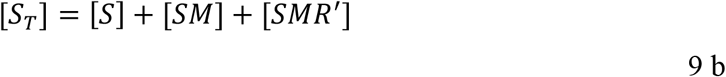

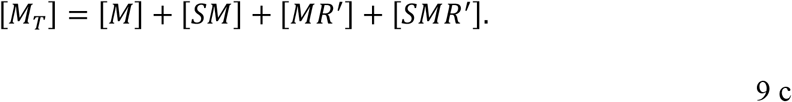

Further I assume that the modulator protein is in detailed balance [22] therefore

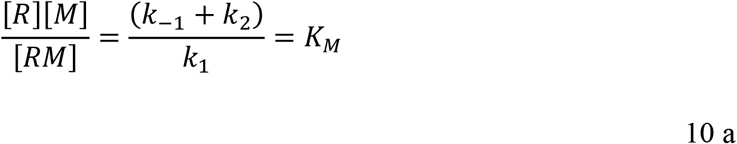

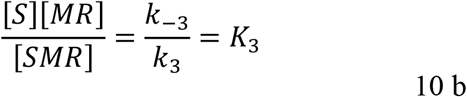

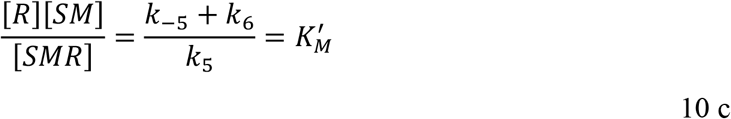

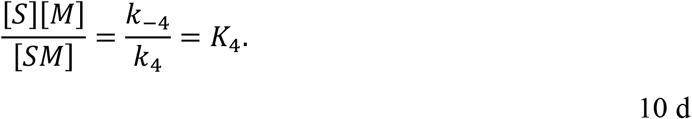

and the dissociation constants need to fulfill:

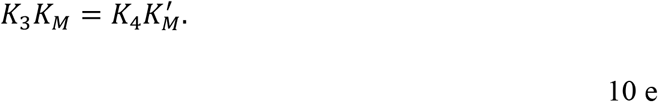

Since protein-protein interactions are fast compared to other cellular processes I assume that the regulator-modulator-signal system is in steady state. This assumption needs to be checked for each experimental system.

### 2.3 Effective parameters of the SAMP-FRET model

One can identify several effective parameters that characterize the SAMP-FRET model. There are two effective allosteric parameters, enzymatic allostery:

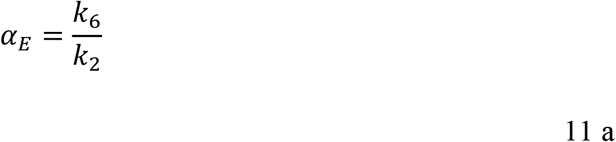

and cooperativity:

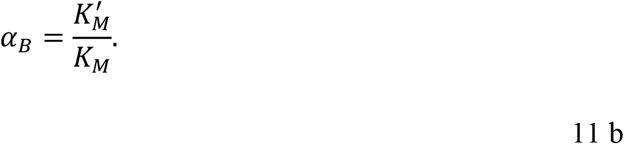

There are three effective parameters that characterize the relationship between the signal-transduction pathway and cellular parameter with the modulator system. The effective regulator affinity given by:

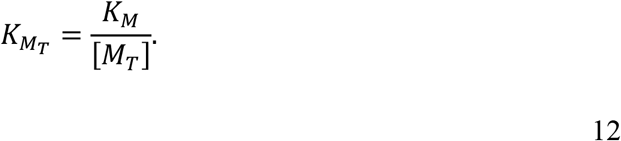

The relative regulator activation rate is given by:

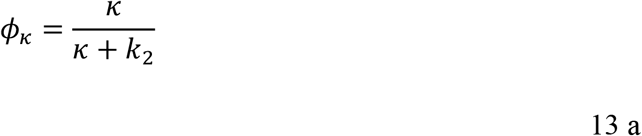

And relative inactivation rate is given by:

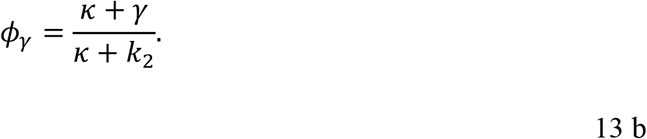

There is one effective parameter that only depends on cellular factors, i.e. the relative modulator concentration:

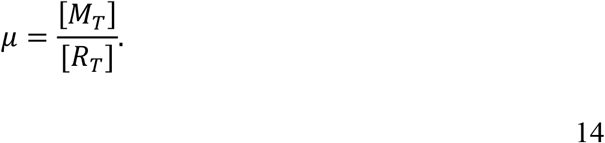

## 3 Results

### 3.1 Qualitative response behavior of the FRET-SAMP sensor depends on the allosteric configuration and the molecular FRET efficiencies

With the model it is possible to systematically investigate the role of allosteric configuration and cellular parameters on the FRET signal. To test how the allosteric configuration influences the FRET response different dose-response curves were simulated with different allosteric parameters. Here model parameters were chosen such that the physiological response to stimulation is maximized [19]. Since also the molecular FRET efficiencies ε_*T*_ and ε_B_ influence the absolute FRET signal they were varied as well (Figure 3).

Figure 4 shows that qualitatively different responses to the signal stimulus are possible. With all three allosteric configurations the signal can lead to both increase or decrease of the measured FRET signal. Here the qualitative response behavior depends on the FRET efficiency of the trimolecular SMR complex. If the orientation of the fluorophores in the trimolecular complex reduces the molecular FRET-efficiency ε_T_ compared to the bimolecular complex by half, the FRET efficiency is reduced with increasing signal concentration. Are the fluorophores in the trimolecular complex oriented such that the molecular FRET efficiency ε_T_ is doubled compared to the bimolecular FRET efficiency ε_B_, the measured FRET value increases with increasing signal concentration.

**Figure 4:**
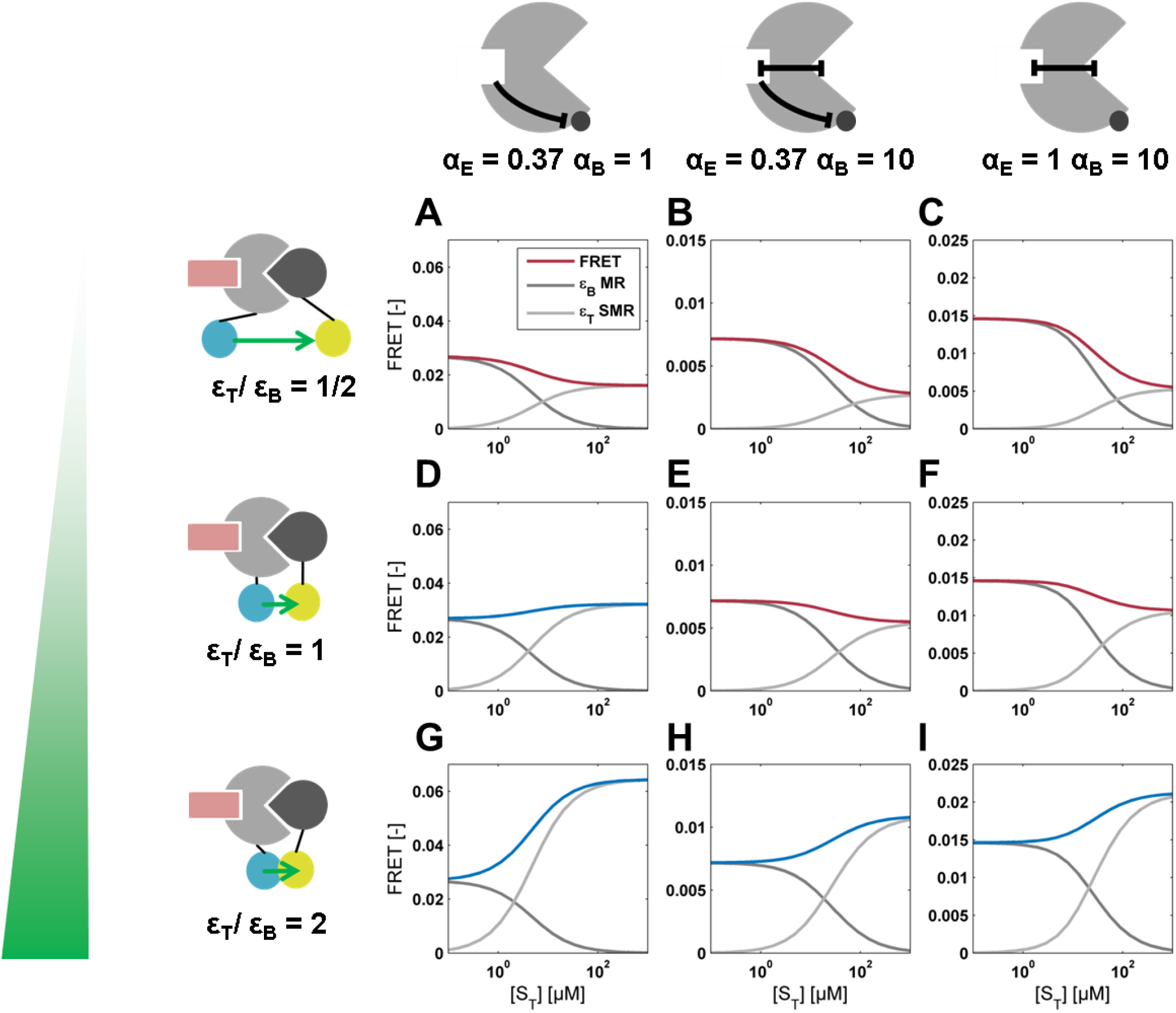
**Orientation of the fluorophore in the trimolecular complex and allosteric configuration influence the qualitative system response. Model parameters in all configurations *ε*_*B*_ =0.1, *k*_*2*_ = 0.7 min^-1^, γ = 10^−4^ s^-1^, *K*_*M*_ = 1 µM, [*R*_*T*_] = 10 µM and *K*_*3*_ = 3.5 µM. For *α*_*E*_ = 0.37 and *α*_*B*_ = 1, [M_T_] = 0.89 µM and κ =0.001 s^-1^. For *α*_*E*_ = 0.37 and *α*_*B*_ = 10, [M_T_] = 0.89 µM and κ =0.002 s^-1^. For *α*_*E*_ = 1 and *α*_*B*_ = 10, [M_T_] = 4 µM and κ =10_4_s^-1^.**

In the case, that the molecular FRET efficiencies do not change, i.e. ε_T_= ε_B_, the qualitative response to stimulation depends on the allosteric configuration. In a SAMP that is only regulated by negative enzymatic allostery, the FRET value increases with increasing stimulus. For a negative coherently regulated SAMP and a SAMP regulated by negative binding allostery, the FRET values decrease with increasing stimulation. In each configuration only the bimolecular MR complex contributes to the FRET signal at low signal concentrations. The concentration of this complex decreases with increasing signal concentration, at the same time the concentration of the trimolecular SMR complex increases. At very high signal concentrations only the SMR complex contributes to the measured FRET value.

How much of the trimolecular complex is formed depends on the allosteric configuration. In a SAMP system that is solely regulated by binding allostery, stimulation with the signal leads to the dissolution of the MR complex so that the fraction of bound regulator is decreased. If both the bimolecular and trimolecular complex have the same molecular FRET efficiency ε this leads to a decreased measured FRET value. For enzymatically regulated SAMPs the MR complex is completely converted to the SMR complex. Due to the slower inactivation rate of SMR complex of the active regulator R’ the concentration of SMR even increases compared to the MR concentration. In a coherent negatively regulated switchable allosteric modulator protein the allosteric modes oppose each other concerning the FRET signal and the qualitative response depends on additional cellular factors.

The simulations show a complex response behavior of SAMP-FRET systems to stimulation. The allosteric configuration and the orientation of the fluorophores in the modulator-regulator and signal-modulator-regulator complex shape this response behavior.

### 3.2 Analytical solution of the model in the case of no or saturating stimulus

To gain more insight into how exactly different parameters determine the FRET response to stimulation, the model is investigated in the limiting case of no signal or a saturating signal concentration ([*S*_*r*_] → ∞). This reduces the models complexity since only the modular-regulator or signal-modulator-regulator complex determine the FRET efficiency.

In the limiting case of no signal, one obtains for the steady-state FRET value the solution:

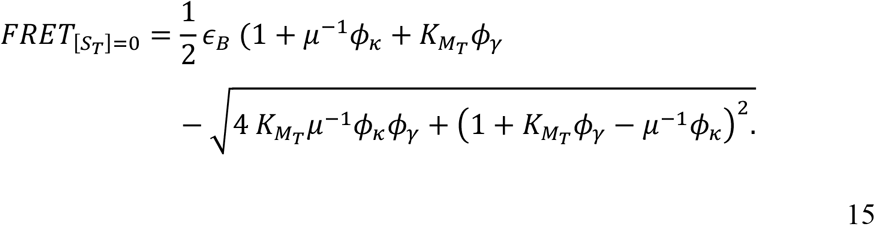

In the limiting case of very large signal concentration, i.e. a saturating stimulus, the FRET efficiency only depends on the SMR complex and one obtains the solution:

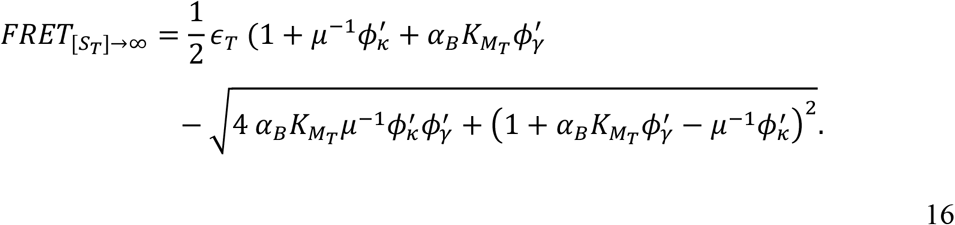

Here the parameters ϕ_κ_ ‘and ϕ_γ_’ do not depend on the inactivation rate k_2_ but on the signal bound inactivation rate *α*_*E*_ *k*_*2*_ = *k*_*6*_:

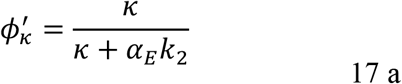

and

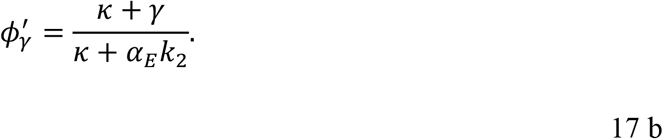

To investigate the interaction between modulator and regulator it is a prerequisite that the measured FRET efficiency is above the detection limit. In addition to the molecular FRET efficiencies and fluorophore choice, the FRET measurement depends on cellular parameters that can systematically be investigated using the analytical solution (Figure 5).

**Figure 5:**
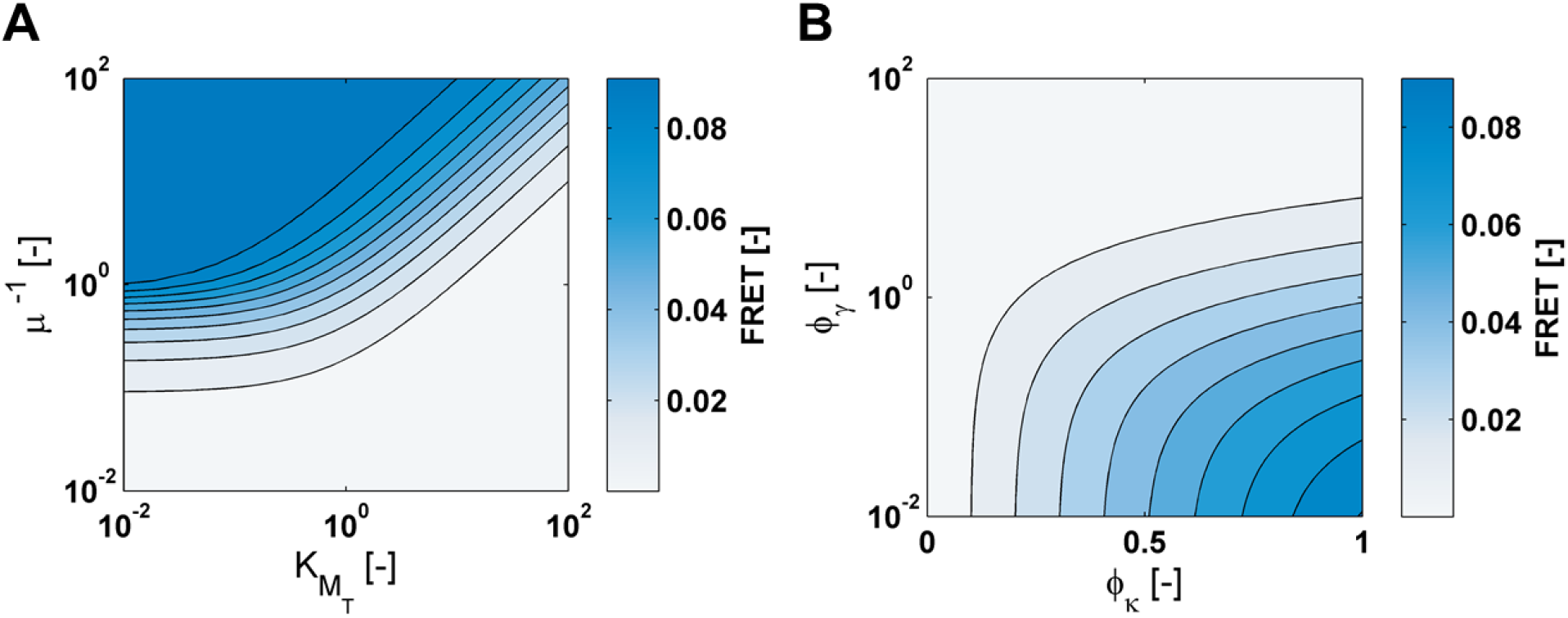
**Influence of the effective parameters on the FRET measurement. (A) FRET as function of *µ*^*-1*^ and *K*_*Mt*_, other parameters *ϕ*_*γ*_ = *ϕ*_*κ*_ = 1. (B) FRET as function of *ϕ*_*γ*_ und *ϕ*_*κ*_, other parameters *µ*^*-1*^ = 1 und *K*_*Mt*_, = 1.**

The effective parameters can be divided into two groups: on one hand parameters that depend on the modulator concentration, on the other hand parameters that depend on the activation rate κ. In the first group belong the relative modulator concentration µ and the effective regulator affinity K_Mt_. When K_Mt_ increases, the affinity to the regulator decreases, therefore it is more difficult for modulator and regulator to form a stable complex. This affects the FRET measurement negatively. If the regulator concentration exceeds the modulator concentration, i.e. µ^-1^ is large, every regulator can bind to a modulator protein and the fraction of bound modulator is large which positively effects the FRET efficiency.

The κ dependent parameters have opposing effects on the FRET efficiency. For large ϕ_κ_ the activation of the regulator exceeds the inactivation by the modulator, increasing the concentration of the modulator regulator complex and so increasing the FRET values. If however the relative inactivation rate ϕ_γ_ is large the unspecific inactivation is larger then the modulator based inactivation. The complex of active regulator with the modulator cannot be formed and no FRET efficiency can be detected.

To detect the interaction of modulator and regulator by FRET *in vivo*, a set of prerequisites need to be fulfilled. This means that the regulator must have a high affinity to the regulator and the regulator concentration should exceed the modulator concentration. In addition, the activation rate of the regulator should be high and the unspecific inactivation of the regulator (i.e. autodephosphorylation) should be low for detectable FRET efficiencies.

### 3.3 Determination of allosteric parameters by variation of cellular parameters

The modulator-regulator FRET sensor should enable the investigation of allosteric signal transduction *in vivo*. It was shown above that the qualitative response to stimulation is not a clear indication of the type of allosteric signal transduction (figure 4).

It was also shown that also cellular parameters influence the FRET efficiency and that those cellular parameters interact with molecular parameters of the modulator, i.e. affinities and enzymatic rates, in the effective parameters. It might therefore be possible that a systematic variation of cellular parameters could uncover the molecular and allosteric parameters of the signal transduction pathway.

Similar to biochemical enzyme kinetic experiments where the variation of the substrate concentration helps identify substrate affinity and enzymatic rates (molecular parameters), variation of cellular parameters could determine those parameters *in vivo*. To this end the influence of cellular parameters is studied in detail.

#### 3.3.1 Half-maximal labeled protein concentration depends mainly on the binding-allostery

It is possible in a cellular context to vary both modulator and regulator concentration at the same time in the same direction. Experimentally this could be achieved by expression of fusion proteins in a bicistronic manner [21]. If the modulator and regulator concentrations are then varied with a constant ratio, this would only change the effective regulator affinity K_Mt_ but not the relative modulator concentration µ.

If no fusion proteins are present, no energy transfer can occur. If the fusion protein concentration is very large the FRET efficiency reaches a maximal value given by:

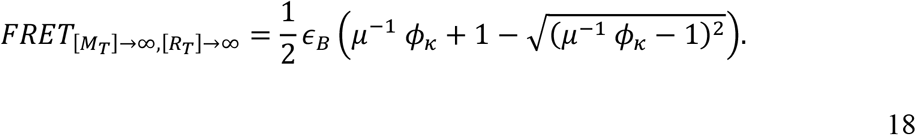

In this case the maximal FRET value depends on the relative modulator concentration µ, the relative activation rate *ϕ*_*κ*_ and is limited by the molecular FRET efficiency *ε*_*B*_. The half-maximal labeled protein concentration is then given by:

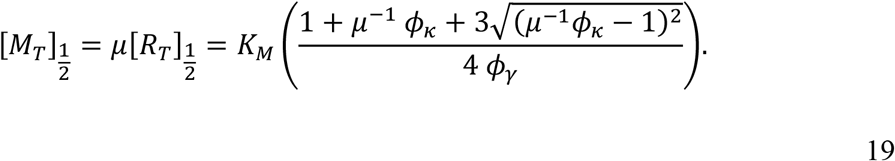

Similar to an experiment for enzyme kinetics the half-maximal protein concentration is mainly determined by a molecular parameter, the Michaelis-Menten constant K_M_. In addition there is a correction factor that includes the relative modulator concentration, and relative activation and inactivation rates.

In the case that the regulator concentration is equal to the modulator concentration, i.e. µ=1, the half-maximal protein concentration simplifies to:

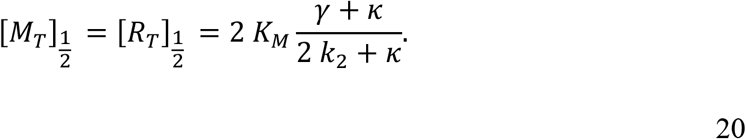

Here the half maximal protein concentration only depends on the enzymatic rate k_2_ if the enzymatic rate is in the same order of magnitude as the activation rate κ. If not, the half-maximal concentration is determined by the Michaelis-Menten constant with a correction factor.

The parallel variation of modulator and regulator concentration could help identify one molecular parameter of the SAMP system, however such an experiment gives no indication about the allosteric signal transduction mode.

To investigate the allosteric mode the same experiment could be performed with a saturating signal concentration. In this case one investigates the molecular properties of the signal-bound modulator. In the case that regulator and modulator have the same concentration and µ=1 the half-maximal protein concentration is given by:

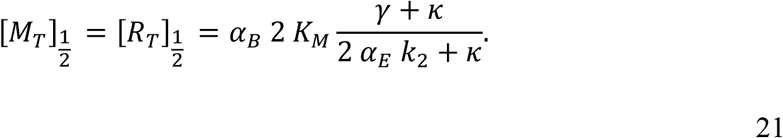

Both enzymatic and binding-allostery change the half-maximal protein concentration under a saturating stimulus. The half-maximal protein concentration directly scales with the binding-allostery parameter α_B_. Experimentally a shift of the half-maximal protein concentration under a saturating stimulus in the amount of the binding-allosteric parameter α_B_ should be observed. The influence of the enzymatic allostery α_E_ depends on the activation rate κ.

As shown above, a saturating signal stimulus can change the measured FRET efficiency independent of the allosteric configuration because of changing molecular FRET efficiencies ε. The half-maximal fusion protein concentration however does not depend on the molecular FRET efficiency and shifts in the half-maximal protein concentration with and without stimulus can directly be related to the binding allosteric parameter α_B_. Possible experimental scenarios with different allosteric configurations are simulated in figure 6.

**Figure 6:**
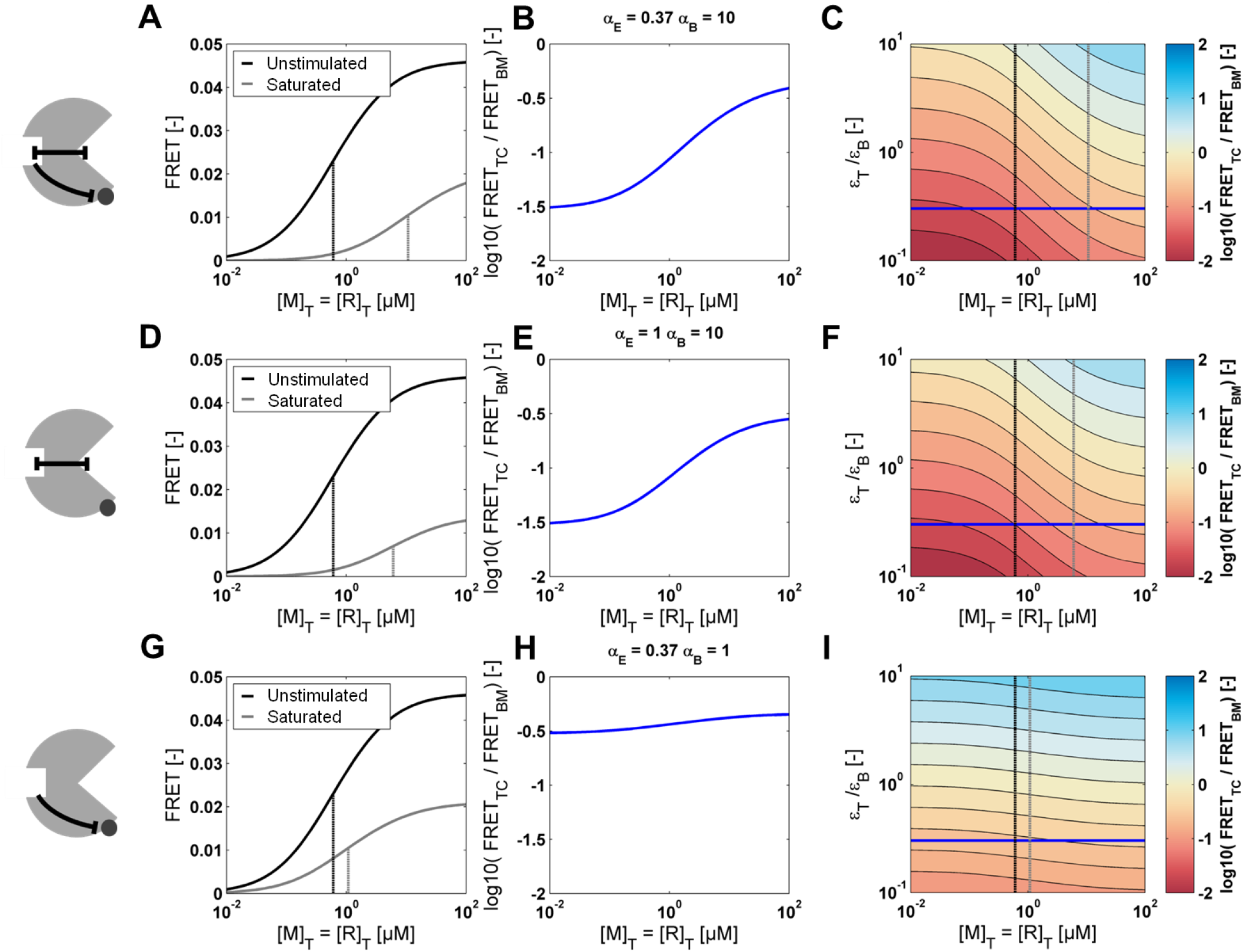
**Influence of the fusion protein concentration on the FRET measurement with and without saturating stimulus. Both regulator and modulator concentration were set to the same value to keep µ=1. (A), (D) and (G): FRET efficiency with and without saturating stimulus. (B), (E) and (H): logarithm of ratio between unstimulated and saturated response curve. (C), (F) and (I): logarithm of ratio between unstimulated and saturated response curve for different molecular FRET efficiencies. (A), (B) and (C) Coherent negative SAMP. (D), (E) and (F) SAMP regulated by binding allostery. (G), (H) and (I) SAMP regulated by enzymatic allostery. Additional parameters: *κ*= 0.01 s^-1^, *γ*= 10^−4^ s^-1^, *k*_*2*_= 0.7 min^-1^, *K*_*M*_ = 1 µM, *ε*_*B*_ = 0.1 and *ε*_*T*_ = 0.03.**

The allosteric configuration changes the half-maximal protein concentration under a saturating signal concentration (figure 6 A, D and G). In a coherent negative allosteric modulator the half-maximal fusion protein concentration increases strongly. If the modulator is regulated by binding-allostery the half-maximal fusion protein concentration still increases, and this increase is minimal in an enzymatic allosteric modulator protein. The shift of the dose-response curve is an indicator of binding allostery.

An informative measure for the shift in the dose-response curves is the logarithm of the ratio of the FRET measurements with and without saturating stimulus:

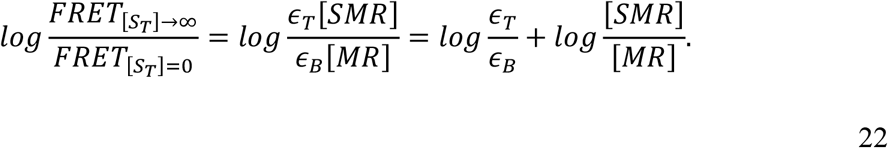

As shown in the middle panels in figure 6 the logarithmic FRET ratio increases with the fusion protein concentration if the SAMP is regulated by binding allostery. It remains constant if the modulator is solely regulated by enzymatic allostery. This effect is independent of the molecular FRET efficiencies of bimolecular and trimolecular complex as shown in the right panels. The contour lines in the right panels are parallel to each other, showing that for each ratio of molecular FRET efficiencies the increase in logarithmic FRET ratio is identical.

#### 3.3.2 Influence of the activation rate κ on the FRET efficiency

It could be possible to manipulate the activation rate κ *in vivo*, e.g. by expression of histidine kinase under control of an inducible promoter system. If no kinase is present, i.e. when κ=0, no regulator is activated and therefore no bimolecular MR complex is formed. For large activation rates *κ* → ∞ the FRET value becomes:

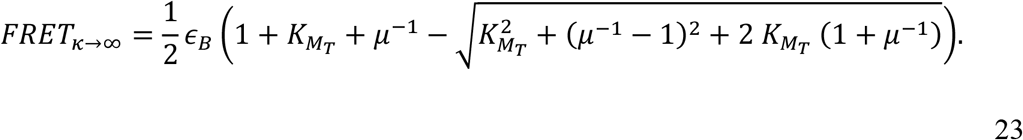

The half-maximal activation rate is then given by:

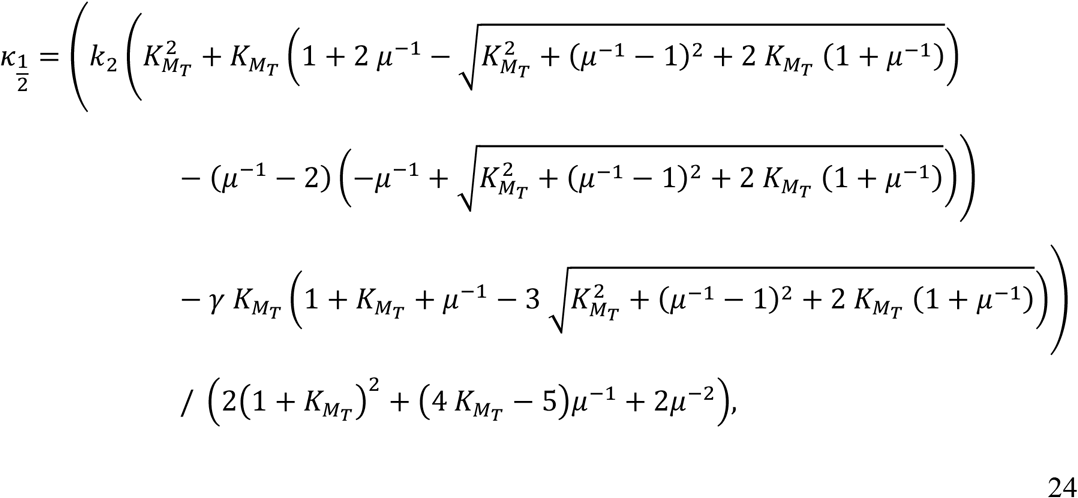

wherein the modulator-based inactivation rate k_2_ and the unspecific inactivation rate γ are single variables and not part of effective parameters. The relative modulator concentration *µ* and the relative regulator affinity *K*_*Mt*_ occur in rational terms. Analogous to the half-maximal fusion protein concentration the half-maximal activation rate is changed by a saturating stimulus. The effect on the dose-response curve is depicted in figure 7.

**Figure 7:**
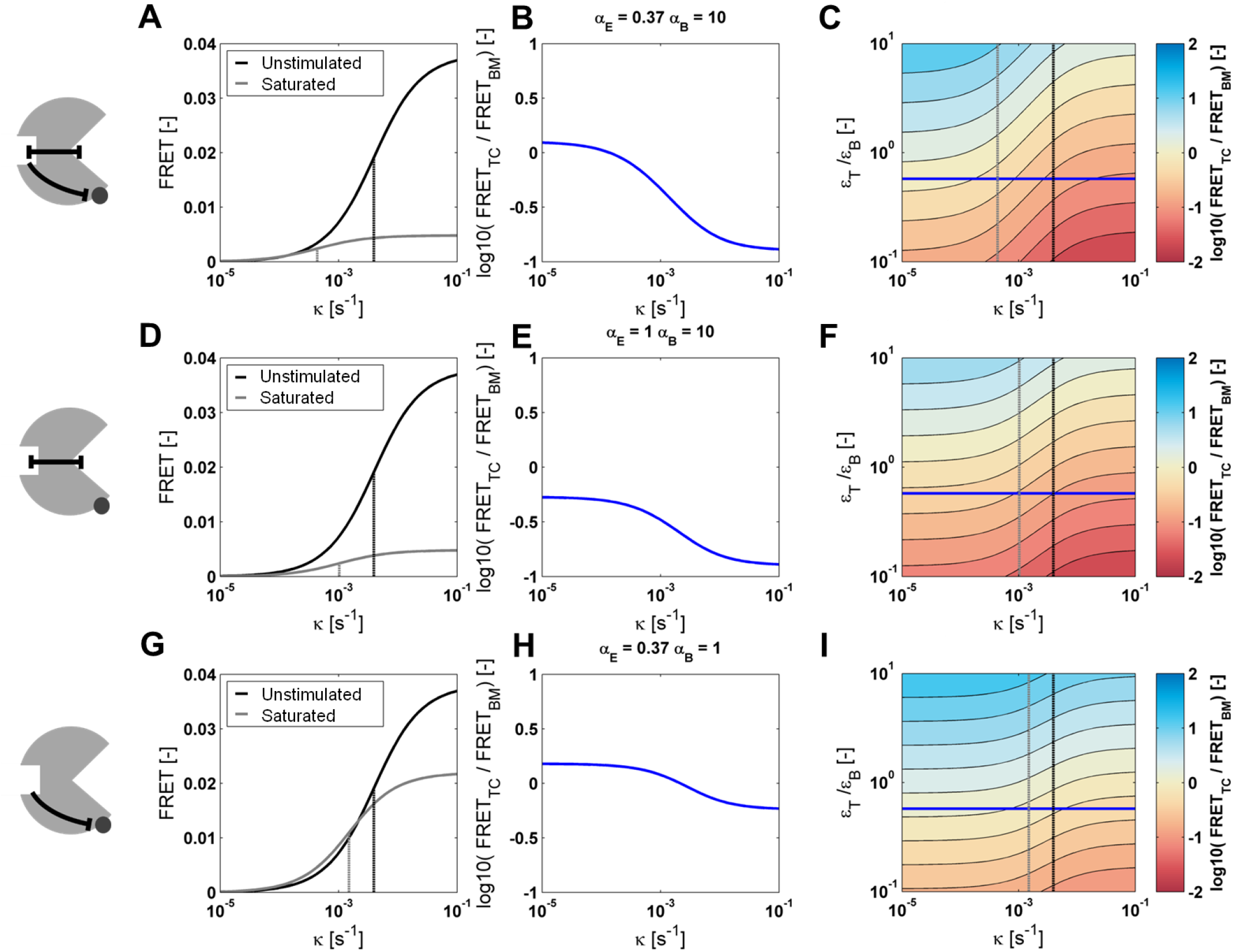
**Influence of the activation rate *κ* on the FRET efficiency with and without saturating stimulus. (A), (D) and (G) show the FRET-efficiency of a unstimulated and stimulated SAMP system. (B), € and (H) show the logarithm of the FRET-ratio of the respective curves. (C), (F) and (I) the logarithmic FRET ratio as a function of different molecular FRET efficiencies(A), (B) and (C) Coherent negative SAMP. (D), (E) and (F) SAMP regulated by binding allostery. (G), (H) and (I) SAMP regulated by enzymatic allostery. Additional parameters: [*M*_*T*_]=[*P*_*T*_] =1 µM, *γ*= 10^−4^ s^-1^, *k*_*2*_ = 0.7 min^-1^, *K*_*M*_ = 1 µM, *ε*_*B*_ = 0.1 and *ε*_*T*_ = 0.03.**

With increasing activation rate, the FRET-efficiency increases. If the system is stimulated with a saturating signal concentration, the FRET-efficiency will decrease at high activation rates. However, at low activation rates in an enzymatically allosteric regulated SAMP and a coherently negative regulated SAMP the FRET efficiency increases with stimulation. The saturating stimulus always shifts the dose-response curve to lower activation rates. This effect is most prominent in a coherently negative regulated SAMP. With SAMPs regulated by binding-allostery or enzymatic allostery this effect is not strong. Because the stimulus decreases the half-maximal activation rate also the logarithmic FRET-ratio decreases with the activation rate. This decrease is again most strong in the coherently negative modulator.

The dose-response curves of the activation rate are changed by stimulation differently depending on the molecular modulator parameters. The shift in the dose-response curve is largest in the coherently negative regulated modulator. SAMPs with binding or enzymatic allostery do not differ in their dose-response curve shift.

#### 3.3.3 Half-maximal relative modulator concentration is influenced in different directions depending on the allosteric mechanisms

When the kinetic of an enzyme is determined, one usually varies the substrate concentration. In a SAMP-system the equivalent experiment would be achieved by varying the regulator concentration (substrate) while keeping the modulator concentration (enzyme) constant. In terms of the effective parameters this would correspond to a change in the relative modulator concentration *µ*. When no regulator is present, no energy transfer can occur. At a saturating regulator concentration the FRET efficiency becomes:

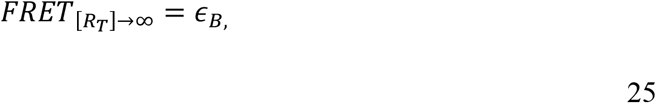

the maximal possible FRET efficiency of an unstimulated modulator, i.e. the molecular FRET efficiency of the modulator regulator complex *ε*_*B*_. The half-maximal regulator concentration then is given by:

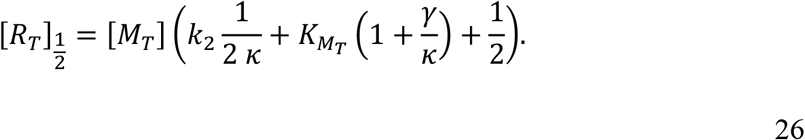

It can be seen that the half-maximal regulator concentration depends on the two molecular parameters, the enzymatic activity *k*_*2*_ and the effective regulator-affinity *K*_*Mt*_. For a large activation rate the half-maximal regulator concentration becomes:

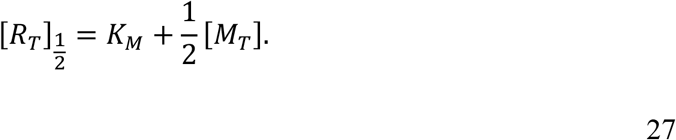

And becomes independent of the enzymatic activity. The saturating signal stimulus changes the half-maximal regulator concentration to:

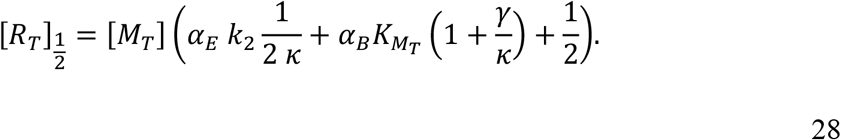

In contrast to changes in kinase-activity and fusion-protein concentration, the saturating stimulus can change the half-maximal regulator concentration in different directions, i.e. increase *or* decrease it depending on the allosteric modes. As shown in Figure 8 D, in a SAMP system regulated by binding-allostery, the half-maximal regulator concentration is shifted to higher concentrations. Enzymatically regulated SAMP-systems shift their half-maximal regulator concentration to lower concentrations upon stimulation (Figure 8 G). In coherently allosteric negative SAMPs the shift in the half-maximal concentration depends on the strength of each allosteric mechanism. In the example shown in Figure 8, the strong binding-allostery leads to a shift to higher regulator concentrations.

**Figure 8:**
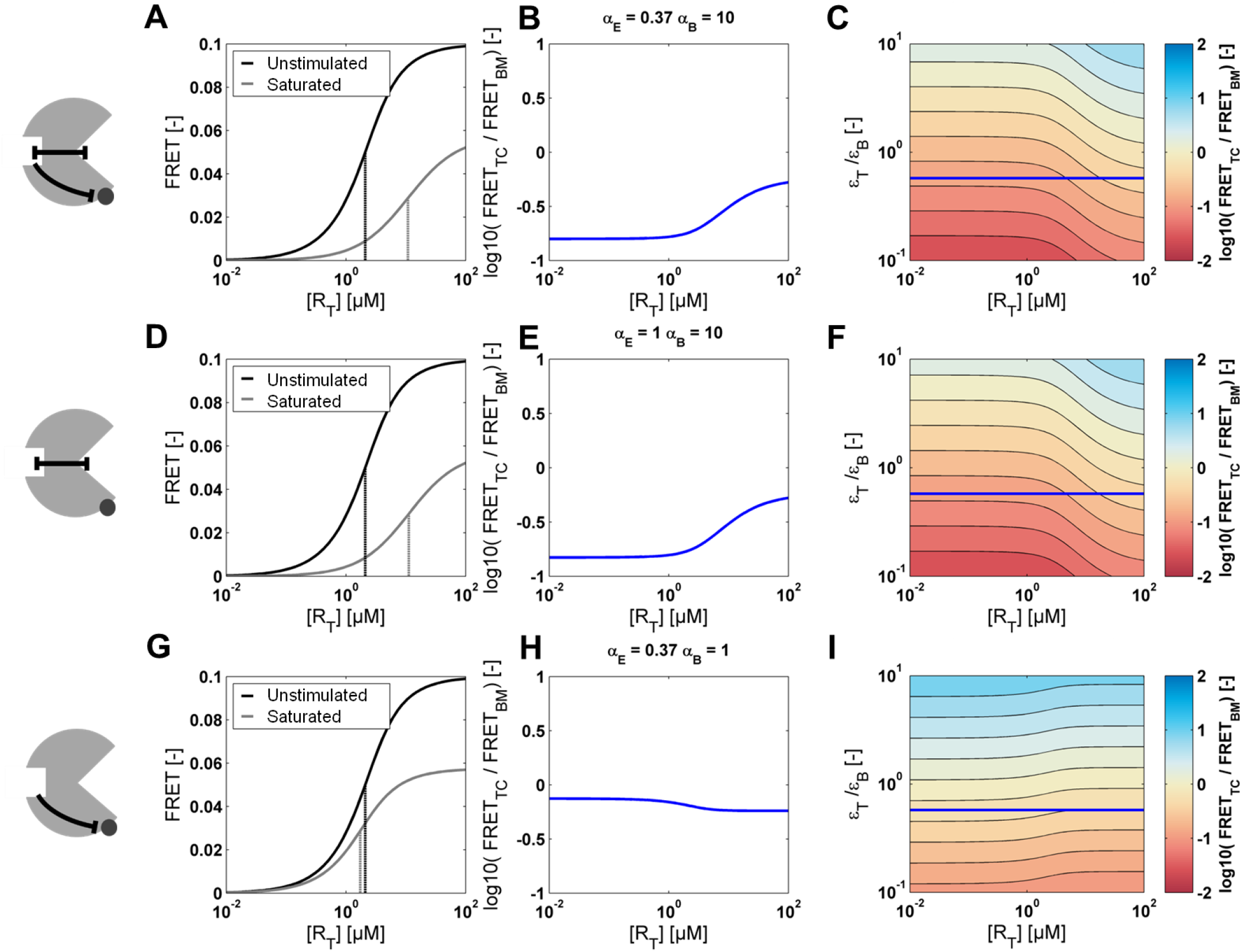
**Influence of the regulator concentration [*R*_*T*_] with and without saturating stimulus. (A), (D) and (G) show the FRET-efficiency of a unstimulated and stimulated SAMP system. (B), € and (H) show the logarithm of the FRET-ratio of the respective curves. (C), (F) and (I) the logarithmic FRET ratio as a function of different molecular FRET efficiencies(A), (B) and (C) Coherent negative SAMP. (D), (E) and (F) SAMP regulated by binding allostery. (G), (H) and (I) SAMP regulated by enzymatic allostery. Additional parameters *κ*= 0.01 s^-1^, [*M*_*T*_]=1 µM, *γ*= 10^−4^ s^-1^, *k*_*2*_ = 0.7 min^-1^, *K*_*M*_ = 1 µM, *ε*_*B*_ = 0.1 and *ε*_*T*_ = 0.06.**

How the dose-response curves are shifted is reflected in the logarithmic FRET-ratio. Enzymatic allostery decreases the FRET-ratio and binding allostery increases it at low activation rates.

For low activation rates, the half-maximal regulator concentration is a qualitative measure for the dominant allosteric mode, i.e. whether a SAMP is regulated by binding or enzymatic allostery. At high activation rates however, the half-maximal regulator concentration only depends on the Michaelis-Menten constant. This is why no influence of enzymatic allostery can be detected anymore.

### 3.4 Discussion

In this paper I present a quantitative model for a bimolecular modulator-regulator FRET sensor. This is the basis for a theoretical understanding of switchable allosteric modulator protein FRET sensors. A necessary prerequisite for the experimental detection of protein-protein interaction is a sufficient activation of the regulator, in addition the regulator concentration should exceed the modulator concentration and the modulator concentration should be larger then the Michaelis-Menten constant of the modulator.

When transient protein-protein interactions are investigated with FRET-reporter systems, often the FRET-efficiency directly correlates with the concentration of the protein-complex of the interaction partners [23,24]. In addition to the concentration of the modulator-regulator complex, in allosteric FRET-sensors, also the conformation of the complex can be influenced by the signal. This can lead to a change in orientation and distance of the fluorophores with respect to each other, which is why the trimolecular SMR-complex can exhibit a different molecular FRET efficiency then the MR-complex at the same concentration.

Formation of the trimolecular complex is a hallmark of allosteric interaction and critical for allosteric signal transduction [25]. The molecular properties of the trimolecular complex are determined by the allosteric regulation of the modulator by the signal. Theoretically, the molecular properties with and without stimulus can be investigated with the FRET sensor, since at saturating stimulus concentration only the trimolecular SMR complex contributes to the FRET signal. Under experimental conditions it is possible to determine the molecular properties of a signal-bound modulator, if an appropriate saturating concentration of the signal is added to the system.

Inspired by *in vitro* measurements, it was investigated if molecular properties of the SAMP system can be identified by variation of certain cellular parameters. Three experimental scenarios were analyzed in detail.

In the first experimental scenario the concentration of both fusion proteins was varied together. Here, the half-maximal fusion protein concentration is mainly determined by the regulator-modulator affinity. It could be shown, that the half-maximal fusion protein concentration is increased by a saturating stimulus. This shift in the half-maximal concentration is mainly determined by the type of allosteric regulation. With enzymatic allostery only a small shift in the half-maximal concentration is observed. Is the modulator regulated by binding-allostery, then the half-maximal fusion protein concentration is shifted proportional to the binding allostery factor α_B_. The shift in the half-maximal fusion protein concentration is a direct indicator of the amount of binding-allostery.

In prokaryotic systems such an experiment can be realized by expression of both modulator and regulator fusion protein from an operon structure with an inducible promoter system. A high activation rate of the regulator is the prerequisite for the experimental detection of modulator-regulator interaction. Due to the high regulator activation one can neglect the effect of enzymatic allostery α_E_.

In the second experimental scenario the activation rate of the regulator was varied. Here too, the half-maximal activation rate is an indicator for allosteric regulation. The change in the half-maximal activation rate appears to correlate to the general strength of the allosteric regulation. *In vivo* this scenario could be achieved by a system wherein the activator-protein (e.g. a histidine kinase in prokaryotic systems) is regulated by an inducible promoter.

In the third and last scenario the influence of the regulator concentration at a constant modulator concentration was investigated. The half-maximal regulator concentration is influenced by both allosteric modes in qualitatively different manners. Negative enzymatic allostery decreases the half-maximal regulator concentration, whereas negative binding allostery increases it. If, however, the activation rate is large, the effect of enzymatic allosteric regulation on the half maximal regulator concentration is minimized. In this case, only the binding allostery influences the shift in half-maximal regulator concentration. In an *in vivo* setting the modulator fusion protein would need to be expressed constitutively, whereas the expression of the regulator protein would need to be regulated by an inducible promoter.

Especially variation of fusion protein concentration and regulator concentration are informative experiments to determine the molecular properties of a SAMP. If, in addition, a saturating stimulus is applied the shift in half-maximal concentration can directly be correlated to the mode and amount of allosteric regulation. In conclusion, SAMP-FRET reporter systems are theoretically suitable tools to investigate allosteric signal transduction and allosteric modes *in vivo*.

## Abbreviations

CFP: Cyan Fluorescent Protein
EC_50_: Half maximal effective concentration
FRET: Förster resonance energy transfer
HK: histidine kinase
RR: response regulator
RRNPP: Rap, Rgg, NprR, PlcR and PrgX
SAMP: Switchable allosteric modulator protein
TPR: Tetratricopeptide Repeats
YFP: Yellow Fluorescent Protein
Rap: Response regulator aspartate phosphatase

